# TIAM-1 regulates polarized protrusions during dorsal intercalation in the *C. elegans* embryo through both its GEF and N-terminal domains

**DOI:** 10.1101/2023.07.24.550374

**Authors:** Yuyun Zhu, Jeff Hardin

## Abstract

Mediolateral cell intercalation is a morphogenetic strategy used throughout animal development to reshape tissues. Dorsal intercalation in the *C. elegans* embryo involves the mediolateral intercalation of two rows of dorsal epidermal cells to create a single row that straddles the dorsal midline, and so is a simple model to study cell intercalation. Polarized protrusive activity during dorsal intercalation requires the *C. elegans* Rac and RhoG orthologs CED-10 and MIG-2, but how these GTPases are regulated during intercalation has not been thoroughly investigated. In this study, we characterize the role of the Rac-specific guanine nucleotide exchange factor (GEF), TIAM-1, in regulating actin-based protrusive dynamics during dorsal intercalation. We find that TIAM-1 can promote protrusion formation through its canonical GEF function, while its N-terminal domains function to negatively regulate this activity, preventing the generation of ectopic protrusions in intercalating cells. We also show that the guidance receptor UNC-5 inhibits ectopic protrusive activity in dorsal epidermal cells, and that this effect is in part mediated via TIAM-1. These results expand the network of proteins that regulate basolateral protrusive activity during directed cell rearrangement.

**Summary statement:** TIAM-1 activates the Rac pathway to promote protrusion formation via its GEF domain, while its N-terminal domains suppress ectopic protrusions during dorsal intercalation in the *C. elegans* embryo.

## Introduction

During embryonic development, cell fate specification and cellular movements work in tandem to achieve the extensive morphological changes of the zygote into a fully developed embryo. One type of cell rearrangement, convergent extension (CE), is a ubiquitous morphogenetic movement that elongates a tissue along one axis while narrowing it along the orthogonal axis (Huebner and Wallingford, 2018; Walck-Shannon and Hardin, 2014). CE or its variants drive morphogenetic movements during gastrulation (Solnica-Krezel and Sepich, 2012) and neurulation (Williams et al., 2014) and shape organogenesis in tissues such as the vertebrate lateral line and the mammalian cochlea (Driver et al., 2017; Sutherland et al., 2020). Because of its early involvement during embryonic development, failure of CE is often accompanied by severe birth defects, including craniorachischisis and spina bifida (Nikolopoulou et al., 2017).

Many studies of epithelial convergent extension have focused on germband extension in *Drosophila*. During this process cells shorten their anterior-posterior boundaries and extend their dorsoventral boundaries as they undergo oriented neighbor exchange (Bertet et al., 2004; Kong et al., 2017; Pilot and Lecuit, 2005). In contrast to the apical junctional dynamics that appears to drive *Drosophila* germband extension, other examples of epithelial CE rely on basolateral protrusive activity to accomplish cell rearrangement. Examples include the ascidian notochord, the sea urchin archenteron, and the mammalian neural plate (Munro and Odell, 2002a; Munro and Odell, 2002b; Sutherland et al., 2020; Walck-Shannon and Hardin, 2014; Williams et al., 2014). How basolateral protrusive activity in epithelia is controlled during tissue rearrangement is therefore an important question in morphogenesis.

A useful system for identifying novel, conserved, functionally relevant molecular pathways that regulate basolateral protrusive activity during epithelial cell rearrangement is the dorsal epidermis of the *C. elegans* embryo. Dorsal epidermal cells undergo a movement known as dorsal intercalation, in which basolateral protrusions drive mediolateral intercalation (Sun et al., 2017; Walck-Shannon and Hardin, 2014). The WAVE complex is constitutively required for protrusion-triggered intercalation (Patel et al., 2008; Walck-Shannon et al., 2015). Our previous work also showed that the *C. elegans* Rac, CED-10, and the RhoG, MIG-2, promote protrusive activity in dorsal epidermal cells. In their active state, CED-10 and MIG-2 lead to the polymerization and nucleation of actin filaments and subsequent formation of lamellipodia via the WAVE and WASP complexes (Walck-Shannon et al., 2015). We also previously identified an upstream guanine nucleotide exchange factor (GEF), UNC-73/Trio, which acts via its RacGEF domain to regulate Rac/RhoG (Walck-Shannon et al., 2015), but other potential upstream regulators have not been thoroughly investigated.

Here, we establish a role for *C. elegans* TIAM-1, a Rac-specific GEF, in regulating protrusive activity during dorsal intercalation. In *tiam-1* mutants lacking GEF activity, dorsal epidermal cells generate fewer protrusions during intercalation, leading to significantly increased intercalation time. We also identify a novel function for the N-terminal domains of TIAM-1 in inhibiting the formation of ectopic protrusions, which at least in part act independently of the GEF domain. Taken together, our data establish a role for TIAM-1 in regulating protrusive activity during dorsal intercalation through both its GEF and N-terminal domains.

## Results

### *tiam-1* is required for normal temporal progression of dorsal intercalation

Dorsal intercalation is driven by the mediolateral protrusive activity of dorsal epidermal cells. Our previous work identified the *C. elegans* RacGTPase orthologs CED-10 and MIG-2 as partially redundant regulators of protrusion formation via activation of the downstream Arp2/3 activators, WVE-1/WAVE and WSP-1/WASP, respectively. We also showed previously that UNC-73/Trio functions as a RacGEF in promoting protrusion formation (Walck-Shannon et al., 2015). However, given the relatively strong protrusion defects in *ced-10* and *mig-2* null mutants compared with *unc-73* mutants specifically lacking RacGEF activity (Walck-Shannon et al., 2015), we hypothesized that additional RacGEFs may regulate this pathway. There are two other RacGEF orthologs in *C. elegans* that show specificity for CED-10 and MIG-2: VAV-1/VAV1 and TIAM-1/Tiam1 (Demarco et al., 2012; Norman et al., 2005; Vettel et al., 2012). In mammals, the function of both VAV1 and Tiam1 have been extensively studied in the immune system. Disrupted expression of both proteins is tightly associated with uncontrolled cell invasion and metastasis (Bartolomé et al., 2006; Boissier and Huynh-Do, 2014; Gérard et al., 2009; Izumi et al., 2019; Liu et al., 2007; Tybulewicz et al., 2003). Their worm orthologs, VAV-1 and TIAM-1, have been shown to function in axonal growth and guidance (Demarco et al., 2012; Fry et al., 2014; Lin et al., 2022; Malartre et al., 2010; Tang et al., 2019). However, our previous work indicated that knockdown of *vav-1* by RNAi had no effect on dorsal intercalation (Walck-Shannon et al., 2015). We therefore focused on TIAM-1, a multidomain protein including an N-terminal myristoylation domain, EVH1-like domain, PDZ domain, and a GEF domain composed of DH and PH subdomains (Fig. 1A) (Demarco et al., 2012).

**Fig. 1.**
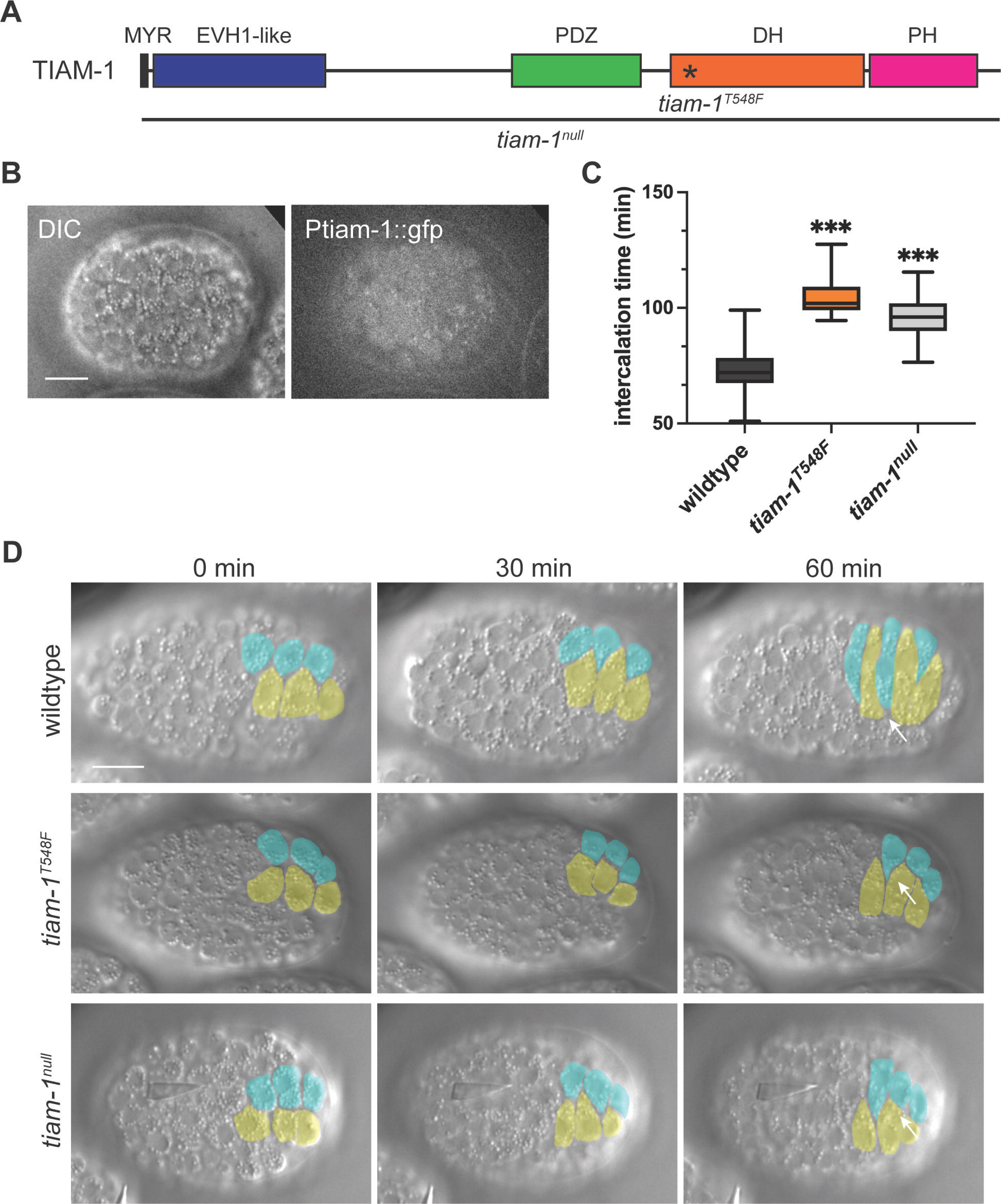
T**I**AM**-1 is required during dorsal intercalation** (A) Protein domain map of *C. elegans* TIAM-1. Domain annotations: MYR, myristoylation site; EVH1-like, Ena/Vasp homology; PDZ; DH, Dbl homology; PH, pleckstrin homology. Asterisk labels the mutation site of allele *tiam-1^T548F^*in the DH domain. Black line labels the deleted region in *tiam-1^null^*. (B) The transcriptional reporter strain *Ptiam-1::gfp* shows that *tiam-1* is transcribed in the epidermis. Left: DIC image, Right: fluorescent image. Scale bar = 10 µm. (C) Intercalation time in *tiam-1^T548F^* and *tiam-1^null^* mutants is significantly longer than in wildtype. ***, p < 0.001, unpaired Student’s T test. (D) DIC images of dorsal intercalation in wild-type (top), *tiam-1^T548F^* (middle), and *tiam-1^null^* (bottom) embryos (dorsal views). Anterior is to the left, posterior to the right; right and left cells are pseudocolored in cyan and yellow, respectively. White arrows point to the leading edge of right-hand dorsal cells at t = 60 min. Cells in wildtype have almost made contact with contralateral seam cells, while cells in mutants have only intercalated halfway. Scale bar = 10 µm.

We first confirmed that *tiam-1* is expressed in the dorsal epidermis. While we could not detect TIAM-1 using an endogenous knock-in strain or via antibody staining of the tagged protein in this line (Y. Zhu and J. Hardin, data not shown), we were able to detect *tiam-1* transcription in dorsal epidermal cells using a *tiam-1* transcriptional reporter strain (Ziel et al., 2009) (Fig. 1B). Embryos from *tiam-1(RNAi)* hermaphrodites displayed delayed completion of dorsal intercalation (data not shown). This defect was validated by quantifying the intercalation time in both a GEF-dead allele, *tiam-1^T548F^* (Tang et al., 2019) and a new *tiam-1^null^* allele we generated, in which the entire *tiam-1* coding region was eliminated using CRISPR/Cas9 genome editing (Fig. 1C,D). These results indicate that *tiam-1* is required for normal dorsal intercalation. Moreover, given the defects displayed by *tiam-1^T548F^*mutants, TIAM-1 appears to function as a RacGEF in regulating dorsal intercalation.

### TIAM-1 activates protrusive activity in dorsal epidermal cells

We next assessed in detail the effects of abrogating the RacGEF activity of TIAM-1 on protrusive behavior of dorsal epidermal cells. In wild-type embryos, dorsal epidermal cells initially generate unpolarized protrusions; shortly thereafter, as cells become wedge shaped, lateral protrusions are retracted and protrusive activity becomes polarized medially (Fig. 2A, left) (Walck-Shannon et al., 2015). We compared this wild-type sequence of behaviors to that displayed by homozygotes for the GEF-dead allele, *tiam-1^T548F^*. Compared to wildtype, *tiam-1^T548F^* mutants showed significantly fewer protrusions during the entire intercalation process (Fig. 2A, quantified in Fig. 2B).

**Fig. 2.**
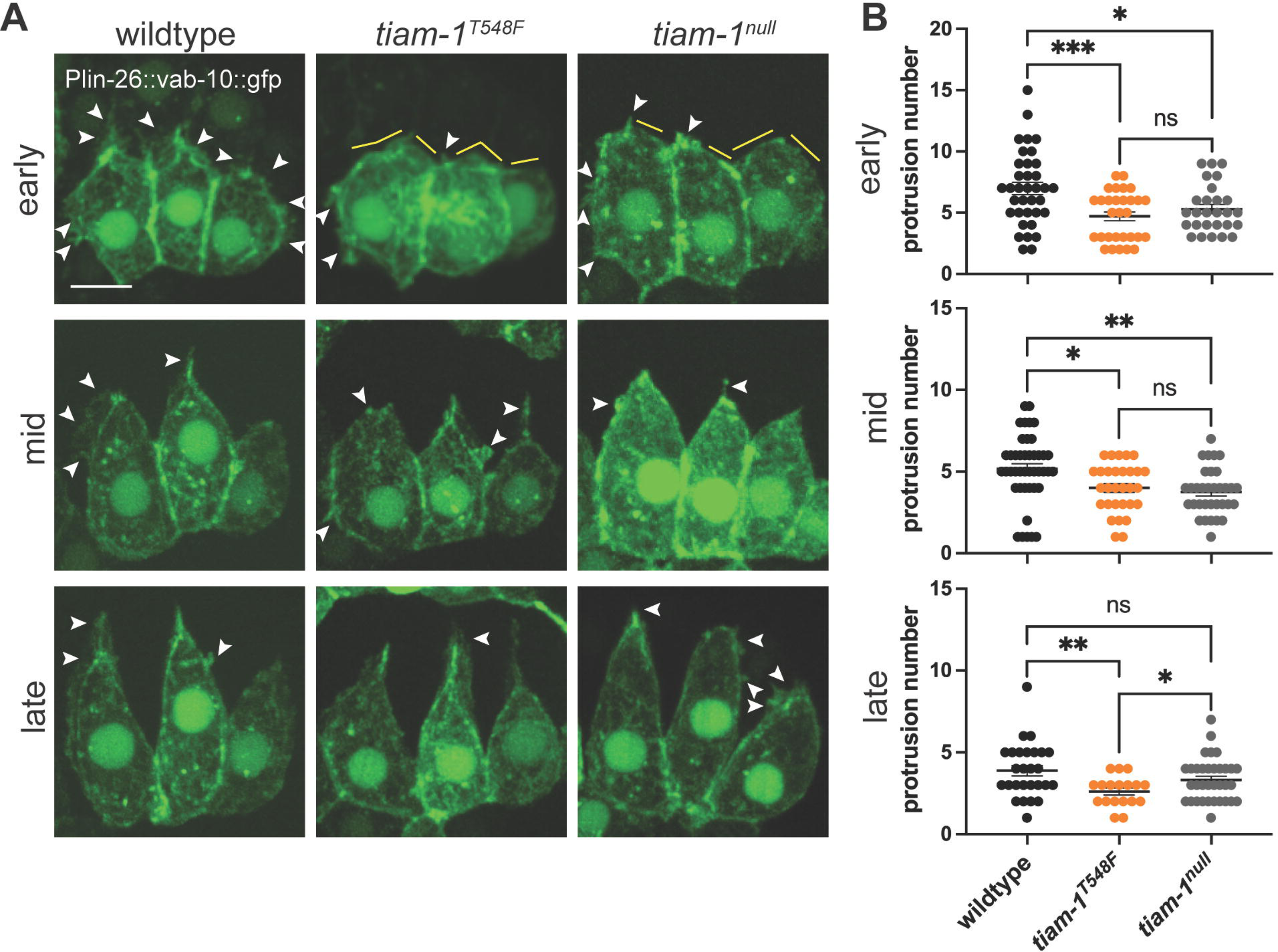
**t*i*am*-1* loss-of-function mutants generate fewer protrusions during dorsal intercalation** (A) Mosaic expression of an epidermal-specific F-actin reporter [*Plin-26::vab-10::gfp*] shows dynamic protrusions during dorsal intercalation in wild-type (left), *tiam-1^T548F^*(middle), and *tiam-1^null^* (right) embryos. White arrowheads indicate protrusions; yellow lines depict the areas lacking protrusions. Scale bar = 5 µm. (B) Quantification of the number of protrusions generated during early, middle and late stages of dorsal intercalation in wild-type and *tiam-1* mutant embryos. Error bars indicate s.e.m. ***, p < 0.001; **, p < 0.01; *, p < 0.05; ns, no significant difference; one-way ANOVA.

We next compared *tiam-1^T548F^* mutants to homozygotes for a true null allele, *tiam-1^null^*. Unexpectedly, while *tiam-1^null^* mutants showed fewer protrusions during the early and middle stages of intercalation, they displayed similar levels of protrusive activity compared to wildtype at later stages, i.e. greater levels of protrusive activity than in *tiam-1^T548F^* GEF-dead mutants, especially later in dorsal intercalation (Fig. 2A, quantified in Fig. 2B). This result suggested that there may be previously unrecognized inhibitory roles for the N-terminus of TIAM-1.

### The N-terminus of TIAM-1 prevents ectopic protrusive activity during dorsal intercalation

Recent work has suggested that the N-terminal domains of TIAM-1 are involved in regulating dendrite development; more specifically, its PDZ domain promotes 4° dendrite branch formation during growth cone morphogenesis (Lin et al., 2022; Tang et al., 2019). We therefore next set out to examine the role of the TIAM-1 N-terminus during dorsal intercalation. Using CRISPR/Cas9, we introduced specific mutations into the *tiam-1* locus, including (1) *tiam-1^ΔN^*, which removes all N-terminal domains, leaving only the GEF domain intact; (2) *tiam-1^ΔPDZ^*; and (3) *tiam-1^ΔPDZ+T548F^*, which introduces the GEF inactivating mutation into the preceding strain (Fig. 3A).

**Fig. 3.**
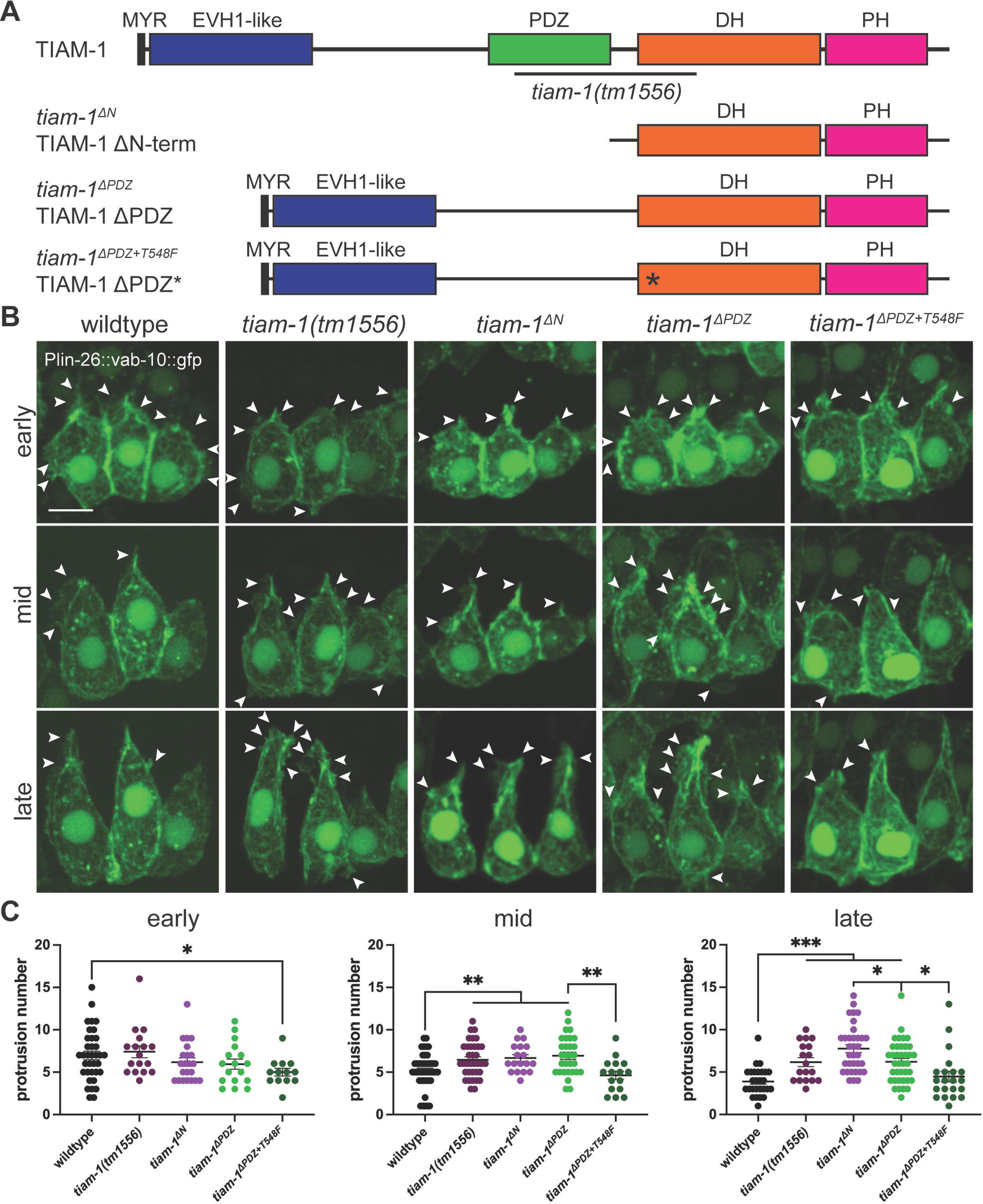
**TIAM-1 N-terminal deletions lead to ectopic and unpolarized protrusions** (A) Domain maps of TIAM-1 and N-terminal deletions relevant to this study. *tiam-1^ΔN^*, truncated protein removing all N-terminal domains, leaving only the GEF domain; *tiam-1^ΔPDZ^*, truncated protein removing only the PDZ domain; *tiam-1^ΔPDZ+T548F^*, truncated protein removing the PDZ domain and containing the T548F point mutation. Black line indicates the lesion in *tiam-1(tm1556)*. (B) Compared to wildtype, *tiam-1* N-terminal deletion mutants display ectopic protrusions, which can be partially ameliorated by eliminating the GEF activity of TIAM-1 (*tiam-1^ΔPDZ+T548F^*). White arrowheads indicate protrusions. Scale bar = 5 µm. (C) Quantification of the number of protrusions generated during early, middle and late stages of dorsal intercalation in wild-type and *tiam-1* N-terminal deletion mutants. Error bars indicate s.e.m. ***, p < 0.001; **, p < 0.01; *, p < 0.05; one-way ANOVA.

We first assessed the intercalation phenotypes of the various mutants using DIC microscopy. Mutants all showed significantly increased intercalation time (Fig. S1A). We next crossed the *tiam-1* mutants with an actin reporter strain to visualize F-actin. Mutant embryos that produce an N-terminally truncated form of TIAM-1 are able to generate significantly more non-polarized protrusions compared to wild-type embryos, specifically during middle to late intercalation, suggesting an inhibitory effect of the N-terminal domains at later stages (Fig. 3B, quantified in Fig. 3C). Compared with *tiam-1^T548F^*, *tiam-1^ΔPDZ+T548F^* mutants generate significantly more protrusions at late stages, whereas they produce significantly fewer protrusions than *tiam-1^ΔPDZ^* mutants (Fig. S1B; Fig. 3B,C), suggesting that the TIAM-1 N-terminus regulates protrusive activity via a previously unknown mechanism that is partly independent of the GEF domain.

### The netrin pathway component UNC-5 is a potential upstream regulator of protrusive activity during dorsal intercalation

TIAM-1 has been implicated in netrin-based signaling in growth cones through its association with the netrin receptor, UNC-40/DCC (Chan et al., 1996; Demarco et al., 2012; Fearon et al., 1990). We hypothesized that netrin signaling might similarly act upstream of TIAM-1 during dorsal intercalation. During neurite outgrowth, the netrin pathway can mediate both attraction and repulsion. In the classic case of *C. elegans* growth cone guidance, attractive signaling acts via the extracellular ligand UNC-6/netrin and UNC-40/DCC. Repulsion often requires the UNC-5 receptor in neurons (Mahadik and Lundquist, 2023; Merz and Culotti, 2000; Norris and Lundquist, 2011; Norris et al., 2014) and other cell types (Su et al., 2000; Ziel and Sherwood, 2010). We therefore examined the localization of UNC-6, UNC-40 and UNC-5 using available knock-in strains (Jayadev et al., 2022).

UNC-6 is expressed predominantly ventrally, without detectable signal in the epidermis near or shortly after the time dorsal intercalation occurs (Fig. S2A). UNC-40 is expressed at a constant level in dorsal epidermal cells throughout dorsal intercalation, with even stronger expression in ventral neuroblasts (Fig. S2B). In contrast, UNC-5 shows strong expression only in the dorsal epidermis (Fig. S2C). UNC-5 is initially visible in cytosolic granules, which we assume are secretory vesicles. As dorsal intercalation progresses, UNC-5 accumulates at the lateral membranes of dorsal epidermal cells (Fig. 4C). Strikingly, UNC-5 showed an increased accumulation at the lateral edges of dorsal cells (Fig. 4B). The junctional localization pattern of UNC-5 makes it a promising upstream candidate involved in the novel inhibitory regulatory function of TIAM-1, and so we focused on it for further analysis.

**Fig. 4.**
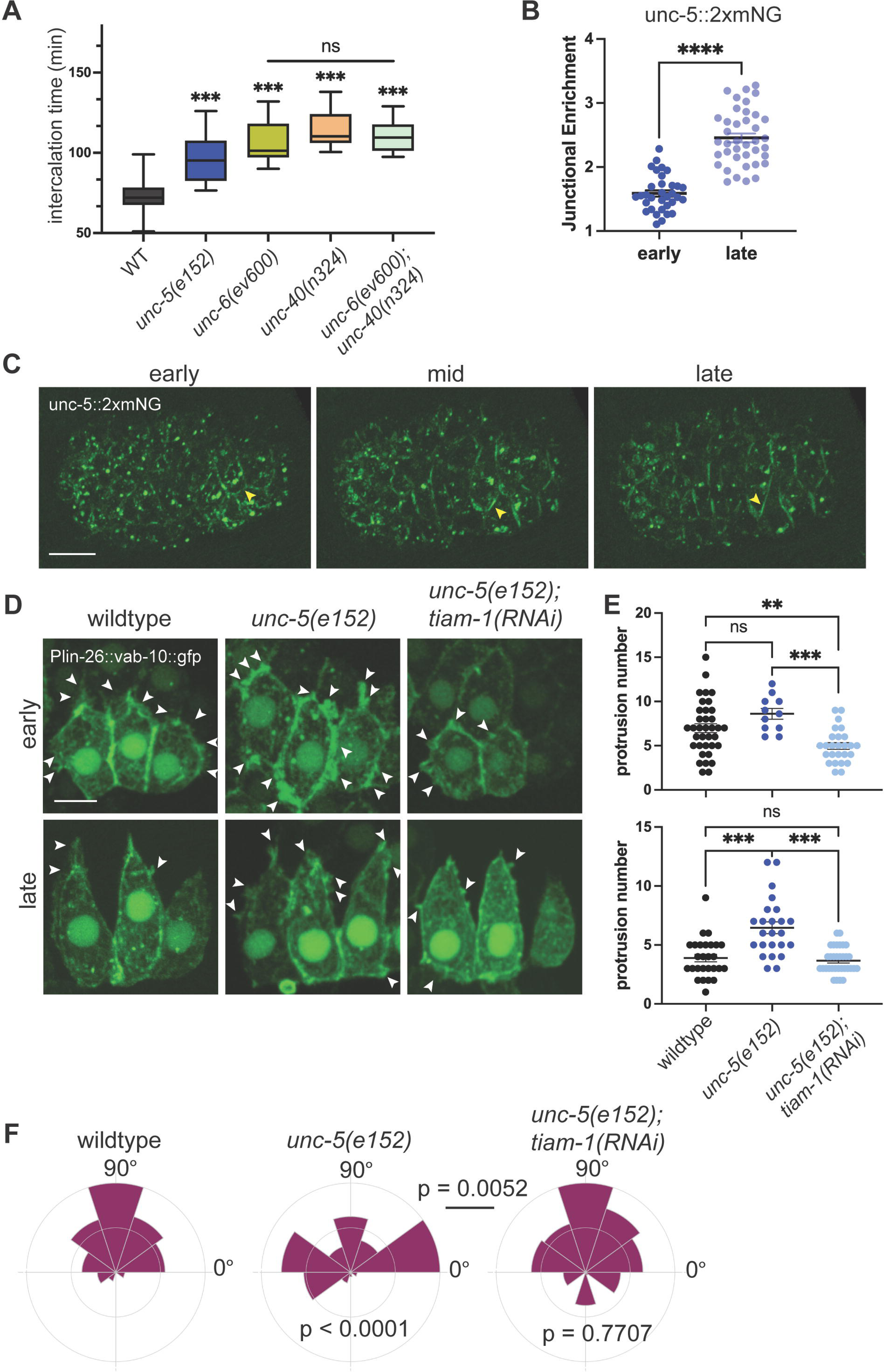
**UNC-5 depletion leads to ectopic protrusive activity** (A) Mutants in netrin pathway components *unc-5*, *unc-6* and *unc-40* display delayed completion of dorsal intercalation. ***, p < 0.001; ns, no significant difference; unpaired Student’s T test. (B) UNC-5 becomes junctionally enriched as dorsal intercalation progresses. ****, p < 0.0001; unpaired Student’s T test. (C) Expression pattern of UNC-5::2xmNG during dorsal intercalation. Yellow arrowheads indicate lateral boundaries of intercalating dorsal cells. Scale bar = 10 µm. (D) Dorsal epidermal cells in a nonsense mutant (*unc-5(e152)*) exhibit ectopic protrusions during late intercalation that are partially suppressed via *tiam-1* knockdown. White arrowheads indicate protrusions. Scale bar = 5 µm. (E) Quantification of the number of protrusions generated during early, middle and late stages of dorsal intercalation in wildtype and *unc-5(e152)* mutants. Error bars indicate s.e.m. ***, p < 0.001; **, p < 0.01; ns, no significant difference; one-way ANOVA. (F) Protrusions show reduced medial polarization in *unc-5(e152)* mutants, which can be partially rescued by RNAi knockdown of *tiam-1*. Rose plots of the angle of protrusion relative to the cell centroid at the late stage of dorsal intercalation. Annotated p values were calculated using the Mardia-Watson-Wheeler test. p values below the plots compare mutants with wildtype; p value above the black line compares *unc-5(e152)* with *unc-5(e152); tiam-1(RNAi)*.

Our previous work had shown that dorsal epidermal cells quickly become polarized at the onset of dorsal intercalation (Walck-Shannon et al., 2015). Since the presence of UNC-5 is often associated with growth cone repulsion (Keleman and Dickson, 2001; Mahadik and Lundquist, 2023; Norris and Lundquist, 2011), the localization of UNC-5 in dorsal epidermal cells suggests a simple model: UNC-5 inhibits protrusive at membrane surfaces where it is expressed, biasing protrusion towards the leading edge. Consistent with this model, dorsal epidermal cells in embryos homozygous for a nonsense allele of *unc-5(e152)* exhibited ectopic protrusions (Fig. 4D, quantified in Fig. 4E) that largely lacked polarity (Fig. 4F). In contrast, *unc-6(RNAi)* and *unc-40(RNAi)* embryos showed no defects in protrusion formation (Fig. S2D). Several previous studies uncovered UNC-40-independent functions for UNC-5, based on comparison of severity of defects between *unc-5* and *unc-40* mutants (Killeen et al., 2002; MacNeil et al., 2009; Mahadik and Lundquist, 2023). These results, combined with our analysis, suggest that UNC-5 likely has an UNC-40-independent function in inhibiting protrusions during dorsal intercalation.

Considering the similarity of defects shown in *unc-5(e152)* and *tiam-1* N-terminal mutants (dorsal cells generate ectopic protrusions and loss of medially polarized protrusions at late stages of intercalation) we next assessed whether there was any connection between UNC-5 and TIAM-1 function during dorsal intercalation. RNAi knockdown of *tiam-1* in *unc-5(e152)* mutants suppressed the ectopic protrusions observed in *unc-5(e152)* mutants alone and partially rescued the polarity defects, suggesting that UNC-5 is involved in regulating Rac-dependent protrusions during dorsal intercalation in part via TIAM-1.

In summary, our results indicate that TIAM-1 acts as a RacGEF in promoting protrusive activity during dorsal intercalation. Its N-terminus also appears to exert a novel inhibitory effect on protrusions independent of its GEF activity, specifically during middle to late stages of intercalation. Finally, UNC-5 may act upstream of TIAM-1 during dorsal intercalation independent of UNC-6 and UNC-40, establishing a novel role for this highly conserved receptor during directed cell rearrangement.

## Materials and Methods

### Strains

*C. elegans* strains were maintained on standard nematode growth medium plates seeded with OP50 *E. coli* at 20°C (Brenner, 1974). Bristol N2 was used as wildtype. Details of strains used in this study can be found in Table S1.

### CRISPR/Cas9 genome editing

All novel mutant alleles with *jc##* designation were generated via plasmid-based CRISPR/Cas9 editing (Dickinson et al., 2015) using repair templates cloned by SapTrap cloning (Schwartz and Jorgensen, 2016). Both deletion mutations (*syb4155* and *syb4244* alleles) were generated by SunyBiotech (Fujian, China). Guides, homology arm primers, and single-stranded repair templates for all CRISPR/Cas9 editing can be found in Table S2.

### RNA interference

Feeding RNAi was performed as described by transferring L4 worms to plates seeded with *E. coli* expressing dsRNA from the relevant target gene or L4440 (control) vectors overnight (Walston et al., 2004). The following day, embryos were dissected from RNAi-treated hermaphrodites and mounted for imaging.

Injection RNAi was performed as described previously (Walston et al., 2004). dsRNA was generated using T7 Megascript kit (Invitrogen). dsRNA was injected at a concentration of 2 μg/µL in nuclease-free water. L4 worms were injected and aged overnight before embryos were dissected from mature adults for imaging. The templates for C11D9.1 (*tiam-1*), F55C7.7 (*unc-73*), T19B4.7 (*unc-40*), F41C6.1 (*unc-6*) and control RNAi were obtained from the Ahringer feeding library (Kamath et al., 2003).

### DIC imaging

Four dimensional DIC movies were collected on either a Nikon Optiphot-2 microscope connected to a QiCAM camera (QImaging) or an Olympus BX50 microscope connected to a Scion CFW-1512M camera (Scion Corp.) using Micro-Manager software (v. 1.4) (Edelstein et al., 2010; Edelstein et al., 2014). All embryos were mounted on 10% agar pads in M9 solution as previously described (Raich et al., 1999).

### Confocal imaging

Embryos were dissected from adult hermaphrodites and mounted onto 10% agar pads in M9 solution and imaged. For fluorescence imaging, a Dragonfly 500 spinning disc confocal microscope (Andor Corp.), mounted on a Leica DMi8 microscope equipped with a Zyla camera and controlled by Fusion software (Andor Corp.), was used to collect images using 0.3 μm slices with a 100x/1.3 NA oil immersion Leica objective at 20°C. Movies were acquired either over a period of 30 minutes at 20 second intervals, or over a period of 60 minutes at 1-minute interval.

### Quantification of dorsal intercalation time

Quantification of intercalation time was performed from DIC movies of embryos mounted on agar pads. For this study, we defined intercalation time as the time between terminal epidermal (Cpaa.a/p) divisions and when cells met contralateral seam cells. At least 20 embryos from at least 8 mounts were analyzed per genotype.

### Quantification of protrusions

Protrusions were quantified from maximum intensity projections of 13 z-stacks of F-actin reporter expression as described above. Protrusion number was obtained by counting aggregations of F-actin reporter signal extending at least 0.2 µm from the cell body. Angles were obtained by comparing the protrusion extension directions relative to the anterior-posterior axis of the embryo in ImageJ. At least 10 cells from at least 5 embryos were analyzed per genotype. We designated cells as “early” if they were in the process of polarizing into a wedge shape, “mid” if the cell tips were <= 2 µm past the dorsal midline, and “late” if the cell tips were <= 2 µm from the contralateral seam cells.

### Statistical analysis

Data from control and experimental groups were compared using one-way ANOVA with Tukey post hoc testing to assess significance between individual groups using Prism (GraphPad Corp.). Protrusion angle related data were compared using the Mardia-Watson-Wheeler test in R using the Circular package.

## Acknowledgements

Strain *tiam-1^T548F^* was provided by the Bülow lab. Some strains were provided by the Caenorhabditis Genetics Center, which is funded by the National Institutes of Health Office of Research Infrastructure Programs (P40 OD010440).

## Author contributions

Conceptualization: Y.Z., J.H.; Investigation: Y.Z., J.H.; Writing – original draft: Y.Z.; Writing – review & editing: Y.Z., J.H.; Supervision: J.H.; Funding acquisition: J.H.

## Data availability

All relevant data can be found within the article and its supplementary information.

